# Transgenic Cowpea Expressing Synthetic *Bt*Cry1Ab Confers High Resistance to Legume Pod Borer (*Maruca vitrata*)

**DOI:** 10.1101/2025.08.16.670672

**Authors:** Muthuvel Jothi, Sanjeev Kumar, Devendra Kumar Maravi, Illimar Altosaar, Vinay Kalia, Lingaraj Sahoo

**Author notes:** Email ID of the corresponding author.

## Abstract

Cowpea, an important food legume crop, suffers significant yield losses due to insect pests, particularly the legume pod borer (*Maruca vitrata*). Narrow genetic base and lack of sufficient resistance to *M. vitrata* in existing cowpea germplasm, make them less effective in managing *M. vitrata* populations. To address this challenge, a transgenic approach is explored. In this study, we develop transgenic cowpea with enhanced resistance to *M. vitrata* by overexpressing a synthetic *cry1Ab* gene, driven by constitutive CaMV35S viral promoter, in an Indian cowpea cultivar. High expression of Cry1Ab was detected in leaves and pods. Bioassays with *M. vitrata* and *Helicoverpa armigera* larvae demonstrated significant resistance of transgenic lines to these lepidopteran pests. The transgenic lines exhibited reduced pod damage, decreased larval feeding, and increased insect mortality compared to non-transgenic controls. NMR analysis demonstrated absence of any new metabolites in transgenic plants. Furthermore, these transgenic lines showed no penalty on growth and development. The transgenic lines exhibited no observable phenotypic drawbacks, indicating their promise for sustainable agricultural practices. Our findings suggest that these transgenic cowpea lines could offer broad-spectrum protection against these pests, providing an effective pest management solution.

## Introduction

Cowpea (*Vigna unguiculata L*.) is an important grain legume that serves as a vital source of high-quality dietary protein for millions of people worldwide, particularly in Africa and Asia (Muthuvel et al. 2021). The resilient nature of the crop to drought and poor soil conditions, and ability to improve soil fertility by nitrogen fixation, makes cowpea cultivation profitable and sustainable in low farm input agriculture. Besides rich in protein, its seeds also have abundant number of essential vitamins (e.g., thiamine folic acid, and niacin), iron, minerals, and dietary fiber (Kamara et al. 2010). Therefore, cowpea which serve as an important source of food and nutrition for both humans and livestock (Muthuvel et al. 2021). Cowpea is predominantly cultivated in India, Africa, the Middle East, and South America (FAOSTAT, 2023). However, cowpea production in major consuming countries fall much short from meeting the domestic demand (Al-Hassan and Diao 2007).

The low yield is mainly due to insect pest, Lepidopteran Pod Borer (LPB), *Maruca vitrata* that feeds internally both vegetative and reproductive stages of cowpea (Abudulai et al. 2017; Kusi et al. 2019). It is the most devastating pest in the genus Maruca that infests 73 host plants ranging from wild to cultivated Fabaceae in tropical Asia and sub-Saharan Africa (Srinivasan et al. 2021). Cowpea is one of the major hosts for *M. vitrata*, with yield losses of up to 72% in Asia and Africa (Srinivasan et al. 2021). Due to its rapid growth, the pod borer exhibits strong adaptability in regions with high humidity, posing significant challenges for effective control. In field conditions, pod borer infestation begins at 25–30 days after the early vegetative stage inflicting maximum damage to flowering stage in cowpea. Young larvae form webs on the reproductive structures and bore into buds and pods to feed on developing seeds (Srinivasan et al. 2021). The application of insecticides is the prominent method to control *M. vitrata* infestation. However, these expensive insecticides are far from reach to marginal farmers in these countries. Over-reliance on synthetic pesticides for the management of LPBs negatively impacts the beneficial insect pollinators.

Developing improved cowpea varieties resistant to legume pod borer is the only environmentally friendly option to control LPBs, reduce pesticide use, and stabilize the production. Conventional breeding to develop LPBs resistant cowpea is limited by absence of functional level resistance in cultivated cowpeas, narrow genetic base and strong cross-incompatibility with wild Vigna species known as reservoir of LPBs resistance. Hence, developing legume pod borer resistance through genetic transformation presents the most feasible option to overcome these constraints (Muthuvel et al. 2021).

*Bacillus thuringiensis (Bt)* insecticidal proteins (Cry proteins) are highly specific to their target pests, with Cry1Ab effective against *M. vitrata* Acharjee and Higgins 2021. While few transgenic cowpea lines expressing Cry1Ab proteins have been developed in a cowpea with African genetic background, field-tested and commercially approved, however, the adaptability and applicability of African cowpea genotypes in Asiatic conditions is questionable owing to the vast difference in cultivation seasons, practices, and genetic background of cultivars (Addae et al. 2020). Recent studies, such as the expression of the Cry2Aa protein in Indian cultivar of cowpea, have proven its effectiveness against *M. vitrata* in India (Kumar et al. 2021). Previously, we reported development of transgenic cowpea plants expressing Cry1Ac gene (Bakshi et al. 2011). Incomplete protection against target pests has been reported in some cases due to reduced Bt toxin expression, late-season declines in efficacy, or variation in expression across plant organs(Jadhav et al. 2020). Such reduced effectiveness during commercial production may heighten the risk of resistant insect populations. To address this, enhancing gene expression—either through the use of strong promoters or codon optimization—offers a promising strategy for conferring complete resistance against *M. vitrata* larvae.

In this study, we have developed transgenic cowpea with a synthetic *crylAb* gene for resistance against the legume pod borer M. vitrata. The *Btcry1Ab* gene, driven by a *CaMV35S* promoter, has shown high expression levels of Cry1Ab at the vegetative and early/late maturity stages of the plant, thereby providing enduring resistance to cowpea. The insect bioassay of Cry1Ab-expressing leaves and immature pods showed minimal damage and high larvae mortality rate compared to non-transgenic control. These transgenic Cry1Ab-expressing cowpea plants could be used for the development of enhanced field-level functional resistance against *M. vitrata* and *H. armigera*.

## Materials and methods

### Construction of binary vector and development of cowpea transgenics

The *Btcry1Ab* construct was kindly provided by Prof. Illimar Altosaar, University of Ottawa, Canada (Shu et al. 2000). The coding sequences for the insecticidal protein Cry1Ab from B. *thuringiensis* were chemically synthesized with codon optimization tailored for plant expression systems (Cheng et al., 1998; Sardana et al., 1996). The full-length plant synthetic *Btcry1Ab* ORF (1.86 kb), with XhoI and KpnI fragment, was cloned into the intermediate vector pRT101 between a 35S promoter and nos terminator sequence. The PstI fragment of 35S::*Btcry1Ab:nos-Ter* was subsequently digested and sub-cloned into binary plant vector pCAMBIA2301. The plasmid construct was then mobilized into disarmed hyper-virulent *Agrobacterium* strain EHA105 and used for cowpea transformation. The T-DNA of binary plasmid pCAMBIA2301 includes neomycin phosphotransferase gene (*nptII*) and β-glucuronidase gene (*gus*) interrupted by catalase intron, both driven by the strong constitutive cauliflower mosaic virus (CaMV) 35S promoter. The cowpea plant transformation was carried out using cotyledonary node explants from an insect-sensitive cultivar cv. PUSA KOMAL (IARI New Delhi) as per our previous protocol (SI Figure 1; Solleti et al. 2008; Bakshi et al. 2011; Kumar et al. 2017). The putative transgenic cowpea plants were established in soil: compost (1:1) and grown to maturity in greenhouse containment. The T_1_, T_2_ and T_3_ seeds were harvested and grown in the soil inside a transgenic poly-house to raise their progeny lines.

### Molecular characterization of transgenic cowpea lines

Genomic DNA was isolated from the young leaves of transgenic plants using the DNASure Plant Mini Kit (Nucleopore, Genetix, India). For screening of the cowpea transformants, PCR analyses were carried out using *cry1Ab* specific primers (Forward: 5’-TGTCCATCTGGTCCCTCTTC-3’ and Reverse: 5’-ATGGTGAAGCCGGTGAGTC-3’) to obtain a partial fragment of 800 bp product from the coding region of the *cry1Ab* gene. A 540 bp fragment of the nptII gene and 1.6 kb of gus gene was also amplified using gene specific primers (*nptII* forward: 5’-GGTGGAGAGGCTATTCGGCTA-3’ and nptII reverse: 5’-GGTAGCCAACGCTATGTCCTGA-3’; *gus* forward: 5’-GGTGGGAAAGCGCGTTACAAG-3’; *gus* reverse: 5’-TGGATTCCGGCATAGTTAAA-3’). The amplified products were resolved by electrophoresis on 1% agarose gel and visualized by ethidium bromide staining (Malke 1990).

Southern blotting was performed using the DIG labeling and detection kit according to the manufacturer’s instructions (Bio-Rad, Hercules, USA). The T_3_ PCR positive transgenic and untransformed cowpea plants were chosen to extract genomic DNA (Maxiprep MN plant DNA maxi kit, Germany). 50-60 µg of total genomic DNA were digested using HindIII, fractionated in 0.8% agarose gel and transferred to a positively charged Zeta-probe membrane. The blot was hybridized with DIG labelled 1 kb PCR product, corresponding to the coding region of *cry1Ab* gene. The hybridized blot was processed for pre and post hybridization wash as per the manufacturer’s instructions. The detection of cry1Ab junction fragment was performed using the instructions of the DIG labeling and detection kit (Roche Diagnostics, Mannheim, Germany).

### Expression analysis of cry1Ab gene in transgenic lines

Total RNA was isolated from the Southern positive plants using a NucleoSpin RNA Plant Kit (TAKARA, Clontech, Japan) and were subjected to cDNA synthesis using RevertAid™ H Minus first-strand cDNA synthesis kit (Fermentas, USA).

Relative fold expression by real-time PCR (RT-PCR) was performed to quantitate the expression of *cry1Ab* gene using specific primers (*cry1Ab* forward: 5’TGG TAC AAC ACT GGC TTG GA3’; *cry1Ab* reverse: 5’-ATGGGATTTGGGTGATTTGA-3’) using USB VeriQuest SYBR Green qPCR Master Mix (2X) (Affymetrix, USA) on a Rotor-Gene Q Real-Time PCR System (Quiagen, Germany). The *VuUBQ1* gene (GenBank Accession No. FG859491) was used as an internal control. The experiment was repeated twice independently with three biological replicates each. The relative expression of *cry1Ab* in wild-type (WT) and transgenic cowpea lines was estimated by using the 2^-ΔΔCt^ method and student’s t-test was used to calculate significance.

### Analysis of Cry1Ab protein in transgenic lines

The expression of the Cry1Ab protein was analyzed in T_3_ transgenic lines generated from four independent transformation events by ELISA and Western blot hybridization. The detection of Cry1Ab protein in leaves and immature pods was assayed by ELISA using antibody coated plates as per the manufacturer’s instructions (Envirologix Quantiplate kit for Cry1Ab/Cry1Ac, Amar Immunodiagnostics, India). Total soluble proteins (TSP) from transgenic and non-transgenic cowpea plants using protein extraction buffer (50 mM Na_2_CO_3_, 100 mM NaCl, 0.05% TritonX-100, 0.05% Tween-20, 2 mM Phenylmethylsulfonyl fluoride and 1 µM leupeptin) and quantified using Bradford assay (Bradford 1976). The TSP were then added to the wells of antibody-coated plates. A secondary antibody Cry1Ab-Enzyme conjugate was then added to each well and the plates were kept for shaking at room temperature for 1 hr. After washing, a colored reaction was observed after adding the substrate in each well and absorbance was measured at 450 nm and comparative histogram was plotted.

For Western blotting, 30 µg of the TSP was fractionated on 12% sodium dodecyl sulfate polyacrylamide gels with (SDS-PAGE) and blotted onto a PVDF membrane by wet transfer (GE Healthcare; Burnette 1981). The rabbit anti-Cry1Ac primary antibody (Amar immune diagnostics, India) in 1:500 dilution and goat anti rabbit conjugated with horseradish peroxidase (Promega) was used for detection. The blot was developed using substrate (SuperSignal West Dura, Thermo Scientific, USA) for 5 min and the reaction was stopped by washing the membrane with sterile distilled water. A single band of ∼67 kDa corresponding to Cry1Ab protein was detected immunologically in all transgenic plants confirming the stable expression of Cry1Ab protein, whereas no such band was observed in the non-transgenic plants.

### Insect bioassays

Maruca pod borer population were reared in an insect growth room at 27 ±⍰2 °C, 16 h light photoperiod and 70 ± 10% relative humidity for optimal growth and mating for in vitro insect bioassays. Insect bioassays were conducted on T_3_ transgenic cowpea lines of 45 days old, where fully expanded mid-canopy leaves and uniformly immature pods were sampled, and second-instar larvae were used to assess the efficacy of cry1Ab expression, as they are more robust and suitable for evaluating sub-lethal effects and realistic field damage (Addae et al 2020). Non-transformed plants served as the negative control. The leaves and immature pods of the non-transgenic and transgenic plants (confirmed by PCR, Southern, and Western blotting) were fed to the larvae separately to evaluate the insecticidal efficacy of the Cry1Ab protein. The ten second instar larvae were released on leaves and pods wrapped with cotton to sustain moisture in the bioassay dishes. Three replicates were carried out for each transgenic/non-transgenic line. In response to the feeding, the mass gain, life cycle and mortality of the larvae and the extent of damage onto the leaves and pods were recorded regularly up to four days after release of the larvae.

### Morpho-physiological analysis of transgenic cowpea lines

The morpho-physiological analysis of the five *cry1Ab* lines were performed at greenhouse conditions. The level of *Cry1Ab* protein in the different progeny plants was determined by Western blot and were grouped into low, medium and high categories. The phenotypic parameters such as plant height, branch number and total number of seeds and pods formed per plant were observed to determine any off-target effects owing to the *cry1Ab* gene expression. To determine the effect of cry1Ab gene expression on seed mass, seed size, the average seed mass of 10 seeds collected from each line was also recorded.

## Metabolomic analysis using NMR

### Metabolite Extraction

Samples were extracted according to Kim et al. 2010 with minor modifications. Briefly, the frozen leaves were ground to fine powder using prechilled mortar and pestle and lyophilized overnight. The extraction was performed with approximately 25 mg of samples and 1 ml of aqueous methanol (80 % v/v) in micro-centrifuge tube. The samples were vortexed for 30 s and then the mixture was further extracted using ultrasonication water bath for 15 min (with 1 min sonication and 30 s pause) using UP200S, Ultrasonic Processor (Hielscher Ultrasonic GmbH, Germany). The extract was centrifuged at 10,000 rpm for 20 min at 4^0^ C. The resulting supernatant was collected in fresh tube and the procedure repeated twice with the remaining pellets. The resulting supernatant from all extraction was combined and concentrated using SpeedVac (Eppendorf AG, Hamburg, Germany) for 3 h. Finally, the dried sample was dissolved in 600 μl of deuterated water (D_2_O) containing 0.05% (w/w) of 3-(trimethylsilyl) propionic-2,2,3,3-d4 acid, sodium salt (TMSP) as an internal standard and transferred into a 5 mm NMR tubes (Sigma-Aldrich, Sant-Louis MO, USA) for further analysis.

### ^1^H-NMR Spectroscopy and spectral data processing

The NMR spectra for all samples were acquired using Bruker Avance III 600 MHz spectrometer with a DCH CryoProbe (BrukerBioSpin GmbH, Rheinstetten, Germany), operating at 600 MHz and 298 K temperature, with 90° pulse width of 12 kHz. One-dimensional ^1^H-NMR spectra with water pre-saturation were acquired using a standard Bruker pulse-acquire sequence during a relaxation delay of 4 s and a mixing time of 100 ms, 32 scans were collected into 64 k data points over a spectral width of 4801.54 Hz and pulse width of 9.46 μs, with an acquisition time of 6.82 s. Water suppression was obtained by pre-saturation sequence present in the routine experiments. The obtained ^1^H-NMR spectrum was manually corrected for phase and baseline using Mestrenova, and the chemical shift was referenced to TMS at 0.000 ppm, to assess the signal quality and to determine the relative proportion of metabolites. The spectral regions of 0.5–10.0 were bucketed into bins (0.04 ppm) using Mestrenova Software (Version 6.0.2–5475; Mestrelab Research, S.L. Spain). Spectral regions 4.84 containing residual signals from water was excluded to remove the residual water by using Mestrenova software. The resulting data was subjected to multivariate analysis (MVDA).

### Multivariate Statistical Analysis for Metabolomic

The dataset was imported into the MetaboAnalyst 5.0 (https://www.metaboanalyst.ca/), and then, the MVDA was performed by PCA and PLS-DA. All the variables were mean centered and log-transformed for multivariate analysis. Furthermore, OPLS-DA was applied to each sample for pairwise comparison between transgenic samples and the control wild type sample using MetaboAnalyst 5.0. The data were statistically analyzed with one-way analysis of variance (ANOVA), and the significant differences were determined using Tukey’s test to evaluate the significant differences between the transgenic samples and the control wild type sample. Differences with p < 0.05 were considered significant.

### Metabolite identification

Metabolite identification was carried out based on the databases such as Biological Magnetic Resonance Data Bank (BMRB; http://www.bmrb.wisc.edu/) and Human Metabolome Database (HMDB; https://hmdb.ca/) and published literature (Maravi et al. 2022). To determine the possible differences in the related metabolic pathways between control and transgenic samples, pathway analysis was carried out using MetaboAnlyst 5.0 and their contributions and biological interpretations were discussed based on the Kyoto Encyclopedia of Genes and Genome (KEGG) database (https://www.genome.jp/kegg/pathway.html).

### Statistical analysis

Statistical analyses were performed using GraphPad Prism, version 8.0 (GraphPad software, USA). The assumptions of normality and homogeneity of variances were evaluated using the Shapiro-Wilk test and Brown-Forsythe and Welch test, respectively. Data met the criteria for parametric analysis, and therefore replicated datasets (n=3 biological replicates) from each experiment were analyzed using one-way analysis of variance (ANOVA) followed by Tukey’s multiple comparison test. The experiment was repeated twice, and a representative image is shown.

## Results

### Development of transgenic cowpea lines with stable cry1Ab integration

Transgenic cowpea plants were produced using *Agrobacterium*-mediated transformation using cotyledonary node explants from the commercial cowpea variety PUSA KOMAL (Kumar et al. 2017). The explants produced an average of 5.4 elongated shoots per explant, with a 92±2% response rate and a maximum transformation efficiency of 3.92% after three selection cycles on kanamycin-supplemented medium (Kumar et al. 2017).

PCR analysis was used to confirm the presence of the nptII, gus, and *cry1Ab* transgenes in the putative T_1_ plants, using gene-specific primers (Fig. 1 A; SI Table 1). Out of 65 plants that survived on the kanamycin selection, 43 tested positives for amplification, yielding fragments of 549 bp, 540 bp, and 798 bp for the *nptII*, gus, and cry1Ab genes, respectively. No amplification products were detected in non-transformed control plants (Fig. 1 B-D). Based on high transgene expression and PCR results, five independent transgenic lines were selected for further analysis to study the segregation patterns of the integrated *cry1Ab, gus*, and *nptII* genes in subsequent generations. For each of these five lines, we analyzed a minimum of 10-12 plants in the T_2_ generation, and at least 8-10 homozygous plants per line in T_3_ generation. Segregation ratios for *cry1Ab, gus*, and *nptII* were recorded in T_1_ and T_2_ generation and were consistent with Mendelian inheritance patterns for a single locus insertion. In T_3_ generation, we selected one homozygous plant per independent line (confirmed by PCR and expression analysis) for insect bioassay and phenotypic evaluation. The segregation ratios of transgenic lines adhered to a Mendelian inheritance pattern (3:1; P < 0.05), indicating that the *cry1Ab* gene was inherited as a single dominant locus (data not shown).

**Figure 1.**
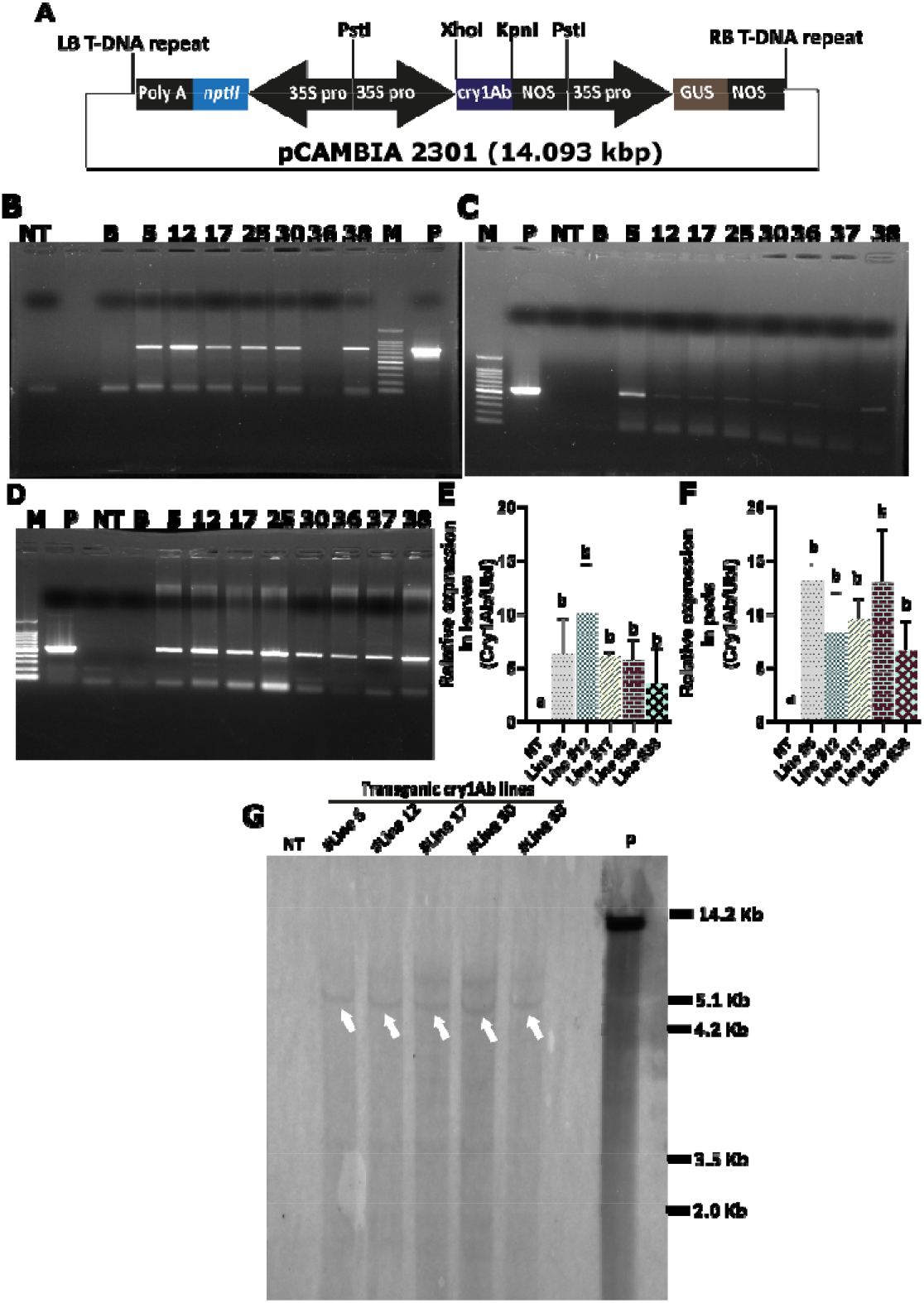
Development and molecular characterization of transgenic *cry1Ab* lines. **(A)** T-DNA region of the *cry1Ab* construct of the binary vector pCAMBIA2301-35Spro::Cry1Ab. (**B-D)** PCR detection of cry1Ab, nptII, and gus using gene specific primers. Lanes, M: 1 kb DNA marker; P: plasmid positive control; NT: non-transgenic cowpea; B: blank as a negative control; Line#5-38: transgenic cry1Ab lines. **(E-F)** RT-PCR based relative fold expression analysis of *cry1Ab* in three independent T_3_ transgenic cowpea lines and non-transgenic (NT) in leaves and pods. VuUbiquitin is used as a housekeeping gene. Bars represent mean values ± standard deviation (SD; error bar) from n=3 biological replicates. Statistical significance was determined using one-way ANOVA followed by Tukey’s multiple comparison test in GraphPad Prism 8.0 Bars shares at least one common letter (a, b) are not significantly different from each other (P> 0.05), while bars labelled with different letters indicate statistically significant differences (P< 0.05) **(H)** Southern blot of HindIII digested genomic DNA of T_3_ transgenic lines (Line #5-38), non-transgenic (NT), and plasmid (P) hybridized with *cry1Ab* probe.

**Table 1.** The metabolite pathway majorly altered in transgenic cowpea pods expressing *cry1Ab*. Total number of pathways and hits along with their negative log values and false discovery rate (FDR), and their impact values are mentioned.

| | Pathway name | Total | Hits | $-\log(p)$ | FDR | Impact |
| --- | --- | --- | --- | --- | --- | --- |
| 1 | Starch and sucrose metabolism | 22 | 7 | 6.6988 | 1.84E-05 | 0.72922 |
| 2 | Alanine, aspartate and glutamate metabolism | 22 | 6 | 5.3342 | 0.000213 | 0.57914 |
| 3 | Galactose metabolism | 27 | 6 | 4.7706 | 0.00052 | 0.36582 |
| 4 | Valine, leucine and isoleucine biosynthesis | 22 | 4 | 2.9363 | 0.026631 | 0 |
| 5 | Cysteine and methionine metabolism | 46 | 5 | 2.5288 | 0.05156 | 0.1793 |
| 6 | Glyoxylate and dicarboxylate metabolism | 29 | 4 | 2.4733 | 0.05156 | 0.27842 |
| 7 | Arginine and proline metabolism | 32 | 4 | 2.3139 | 0.055568 | 0.14435 |
| 8 | Amino sugar and nucleotide sugar metabolism | 52 | 5 | 2.293 | 0.055568 | 0.11106 |
| 9 | Glycine, serine and threonine metabolism | 33 | 4 | 2.2647 | 0.055568 | 0.16478 |
| 10 | Arginine biosynthesis | 18 | 3 | 2.1757 | 0.061389 | 0.23148 |
| 11 | Citrate cycle (TCA cycle) | 20 | 3 | 2.0438 | 0.075611 | 0.19387 |
| 12 | Carbon fixation by Calvin cycle | 21 | 3 | 1.9835 | 0.079632 | 0.05923 |
| 13 | Glutathione metabolism | 26 | 3 | 1.7256 | 0.13312 | 0.05801 |

To verify the expression of *cry1Ab*, relative fold expression by real-time PCR (RT-PCR) was conducted on PCR-positive plants. In the T_3_ lines 17, 30, and 38, expression levels of cry1Ab were found to be 3-to 10-fold higher in leaves and 6-to 13-fold higher in immature pods relative to the *Vuubiquitin* gene. No amplification was observed in control non-transformed plants (Fig. 1 E and F). Expression levels varied significantly between leaves and pods across the transgenic lines. Notably, lines #12 and #5 exhibited the highest *cry1Ab* expression in leaves and immature pods, respectively, while line #38 showed the lowest expression in both tissues among the transgenic lines (Fig. 1 E and F).

The stable integration and copy number of the *cry1Ab* gene were further assessed in the T_3_ lines by Southern blot analysis. HindIII-digested genomic DNA hybridized with a *cry1Ab* probe displayed hybridization bands larger than 4.2 Kbp, confirming the successful integration of the *cry1Ab* gene into the cowpea genome. Southern blotting also confirmed that all the five lines contained a single copy of the *cry1Ab* gene, indicating a single T-DNA insertion event. No hybridization signal was observed in the non-transformed control plants (Fig. 1 G). Three transgenic lines showing similar Southern hybridization patterns were independently generated. The similarity likely reflects preferential integration or conserved restriction site environments.

### Transgenic cowpea lines express higher Cry1Ab protein

The qualitative and quantitative measurement of the Cry1Ab protein expression was analyzed by Western blotting and ELISA, respectively. The total soluble protein from the leaves and immature pods of transgenics and non-transformed control plant was used to test Cry1Ab protein. The Cry1Ab protein that are hybridized with the polyclonal Cry1Ab antibodies was detected as a signal band of molecular weight ∼67 kDa in all tested lines (Fig. 2 A and B). The intensity of the Cry1Ab protein in all the transgenic lines, except #38, was higher in leaves. In immature pods, lines #17 and #38 were showed low Cry1Ab intensity than the rest of the transgenic lines. Faint non-specific bands were detected, likely due to antibody cross-reactivity, but do not interfere with interpretation of Cry1Ab expression. These results on expression of Cry1Ab toxin protein demonstrate stable and consistent expression in most of the transgenic lines. No band was detected in the case of the untransformed control (non-transgenic) plant.

**Figure 2.**
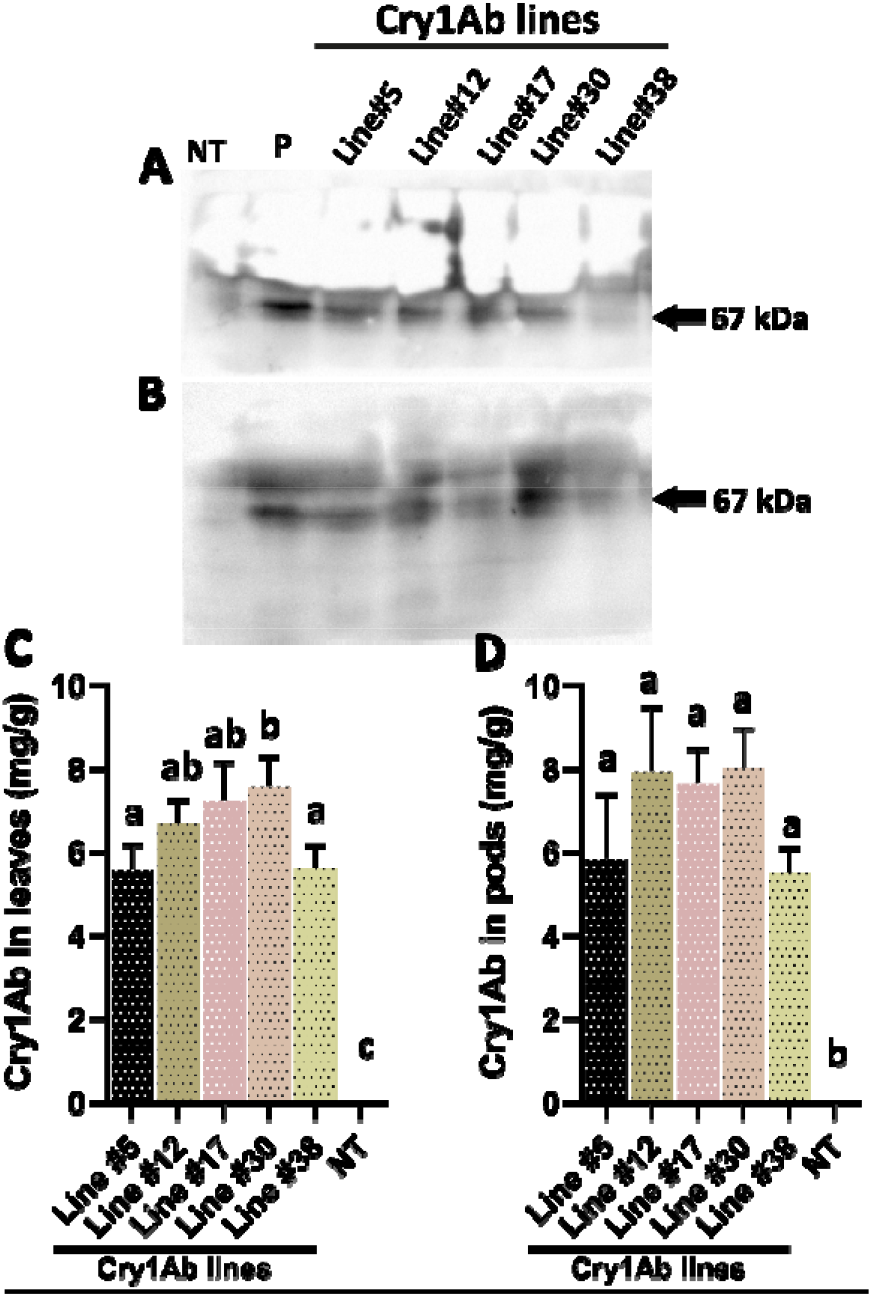
Expression and quantification of Cry1Ab protein in transgenic cowpea. Western blot analysis of *cry1Ab* gene in T_3_ transgenic lines (Line#5-38) and non-transgenic (NT) leaves **(A)** and pods **(B)**. A single band of ∼67 kDa corresponding to Cry1Ab was shown by a black arrow. Purified Cry1Ab protein was used as positive control (P) and proteins from non-transgenic plants were used as a negative (NT). **(C-D)** Quantitative estimation of Cry1Ab protein in T_3_ transgenic lines and non-transgenic (NT) using Cry1Ab-enzyme conjugate. Average quantity of total Cry1Ab protein of the transgenics is represented as mg g^−1^ fresh mass ± standard deviation. Bars represent mean values ± standard deviation (SD; error bar) from n=3 biological replicates. Statistical significance was determined using one-way ANOVA followed by Tukey’s multiple comparison test in GraphPad Prism 8.0. Bars shares at least one common letter (a, b and c) are not significantly different from each other (P> 0.05), while bars labelled with different letters indicate statistically significant differences (P< 0.05).

The concentration of Cry1Ab protein in leaves and immature pods was quantified in five independent transgenic lines using ELISA (Fig. 2 C and D). Since the cry1Ab gene was driven by CaMV35S constitutive promoter, the Cry1Ab protein was expressed in both tissues. We observed that the Cry1Ab protein was highly expressed across the three generations in transgenic cowpea lines. The protein expression ranged from 2.8 to 10.8 mg/g in leaf tissues and 5.0 to 10.7 mg/g in immature pods in T_1_ transgenics (SI Figure 2 A and B). The T_2_ transgenics recorded a maximum level of Cry1Ab protein expression, 2.9 to 10.1 mg/g leaf tissue and 5.6 to 10.1 mg/g young pods (SI Figure 2 C and D). In T_3_ generation, the Cry1Ab protein expression observed to be 5.5 to 7.5 mg/g in leaf tissue and 5.5 to 8 mg/g in immature pods (Fig. 2 C and D). The accumulation of Cry1Ab proteins was higher in the reproductive tissues (young pods) compared to vegetative tissues (leaf) across all the three generations in tested events. Among the five transgenic lines tested, cry1Ab expression level in lines #12, #17 and #30 were found higher, in both vegetative as well as reproductive tissues at T_3_ generation. The concentration of Cry1Ab proteins in transgenic lines were stabilized in subsequent generation. The T_3_ transgenic lines of the respective T_1_ parents, showed 5.7% higher expression (on an average) of Cry1Ab protein in leaves and pods of all transgenic lines.

### Insect bioassay of transgenic cowpea leaves and immature pods showed higher mortality of *M. vitrata* and H. armigera

The entomocidal activity of cowpea transgenic plants, expressing moderate to high levels of BtCry1Ab, was evaluated through insect feeding bioassays. The mortality rate for *M. vitrata* larvae ranged from 33-60%, while H. armigera larvae exhibited a mortality range of 66-96% when fed on leaves of T_3_ transgenic plants (Figure 3). Similarly, in the detached pod assay, mortality rates ranged from 60-90% for *M. vitrata* and 46-63% for H. armigera in the T_3_ transgenic lines (Figure 3). Notably, transgenic lines #30 and #38 showed the highest mortality for both *M. vitrata* and H. armigera larvae in leaf and pod bioassays. Significant differences in mean larval mortality were observed between the transgenic lines and the control. Additionally, surviving larvae on the transgenic plants exhibited retarded growth compared to those on non-transgenic controls (data not shown). For *M. vitrata* larvae, Cry1Ab protein concentrations greater than 6 mg g^−1^ fresh mass in the pods caused over 70% mortality. However, at similar Cry1Ab protein concentrations, H. armigera larvae exhibited relatively lower mortality (over 53%). In contrast, H. armigera larvae showed higher mortality (>83%) than *M. vitrata* in transgenic leaves expressing >6 mg g^−1^ fresh mass Cry1Ab toxin (Fig. 2 C and D; Figure 3). Although larval mortality was positively correlated with Cry1Ab protein levels, the reason for the differences in mortality between *M. vitrata* and H. armigera in leaves and pods remains unclear.

**Figure 3.**
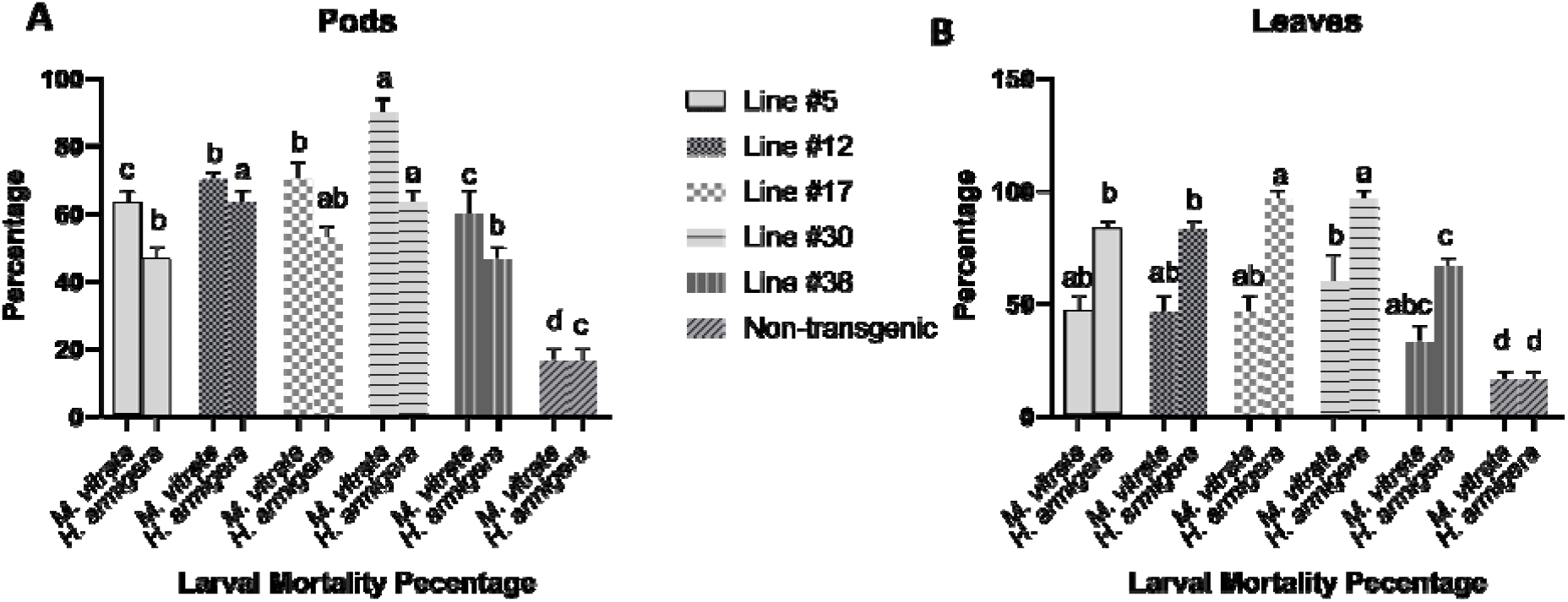
Efficacy of transgenic cowpea lines against *Maruca vitrata* and *Helicoverpa armigera* larvae by detached leaf and pod bioassay from transgenics (Line #5-38) and non-transgenics (NT). Mean percentage values ± standard error (SE) from n=3 biological replicates are provided. Statistical significance was determined using one-way ANOVA followed by Tukey’s multiple comparison test in GraphPad Prism 8.0. Values shares at least one common letter (a, b, c, and d) are not significantly different from each other (P> 0.05), while values labelled with different letters indicate statistically significant differences (P< 0.05).

## NMR

### Metabolite identification

The ^1^H NMR spectrum obtained at 600 MHz for the transgenic and non-transgenic cowpea pods (SI Fig. 3). The NMR results revealed the variation 40 major metabolites non-transgenic cowpea pods. These 40 metabolites include amino acids, organic acids, sugars, and other aromatic compounds (SI Table 2). ^1^H NMR performed on immature pods of transgenic and non-transgenic cowpea, the 600MHz spectra exhibited in three main regions (SI Fig. 3). The first region (0.5-3.0 ppm) contained signals of amino acids and some organic acids were identified. The second region (3.0-5.8 ppm) comprised signals of carbohydrates. Sucrose (5.41 ppm), fructose (4.11 ppm), beta glucose (4.64 ppm), alpha glucose (5.23 ppm) were the most abundant carbohydrates in this region. In the third region (5.8-9.5 ppm), were relatively weak signals which are corresponds to the aromatic groups from phenolic compounds and aromatic amino acids. The most abundant signals in the aromatic region, phenylalanine (7.43 ppm), tryptophan (7.71 ppm), fumarate (6.53 ppm) and formic acid (8.54 ppm) were identified (SI Fig. 3).

### Multivariate statistical analysis for metabolomic

Identification of metabolite variation of transgenic and non-transgenic leaves and pods was carried out by principle component analysis (PCA). In our current study, the first three components explained 78.6% of whole data: PC1, 41; PC2, 19.8; PC3, 17.8%, respectively (Fig. 4). The score plots clearly showed that samples from the transgenics and non-transgenics were well separated (Fig. 4A). However, lines #12 (TR2), #17 (TR3), #30 (TR4), and #38 (TR5) samples showed noticeable overlaps indicating that these samples not be efficiently separated. The control group was negatively influenced by PC1 and positively influenced by PC2, whereas, line #5 (TR1) group was negatively influenced by both PC1 and PC2. The lines #12 (TR2), #17 (TR3), #30 (TR4), and #38 (TR5) groups were negatively influenced by PC1. The loading plots indicated that the discriminative metabolites responsible for the variable separations. Metabolites such as isocitrate and isoleucine are represented in PC1. PC2 was mainly characterized by proline, fructose and malate (Fig. 4B).

**Figure 4:**
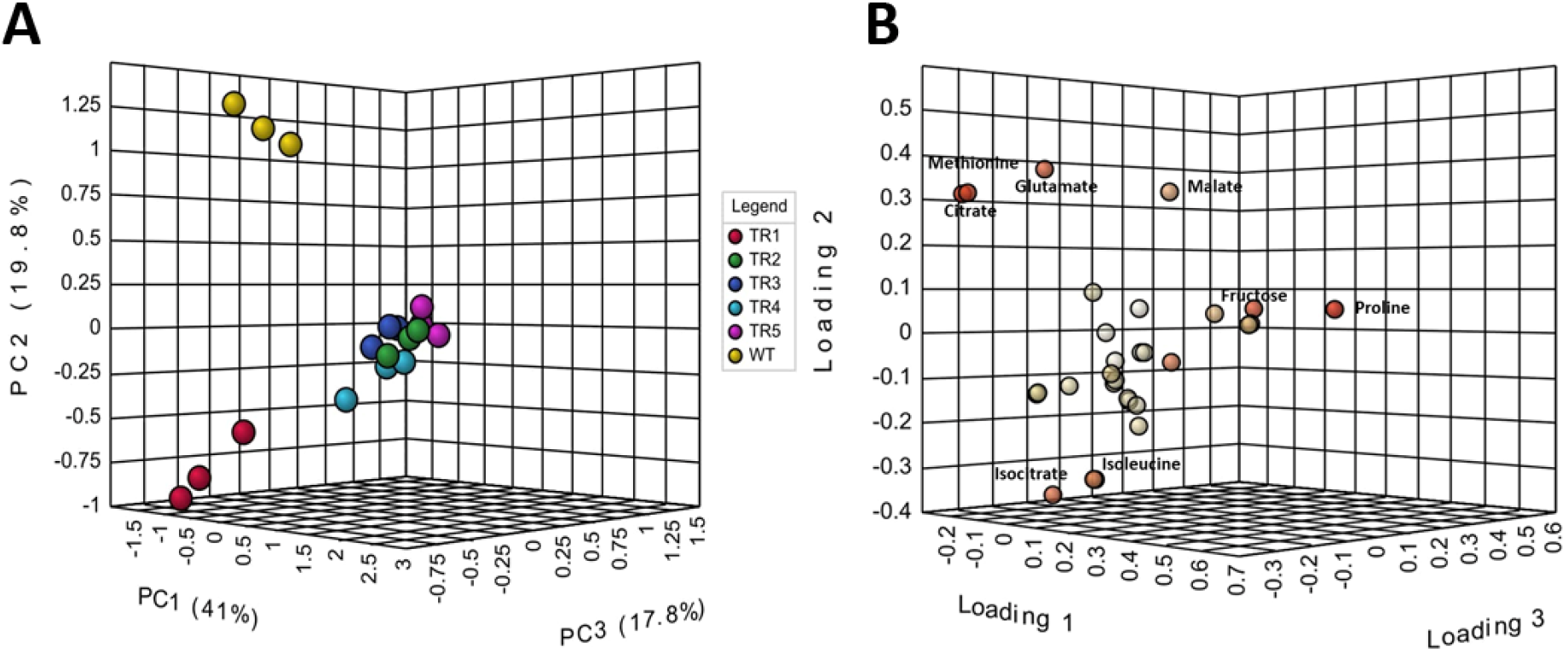
The principal component analysis (PCA) score plot **(A)** and loading plot **(B)** illustrate the ^1^H NMR spectra of transgenic (line #5:TR1, line #12:TR2, line #17:TR3, line #30:TR4, and line #38:TR5) and non-transgenic (WT) cowpea pods. Distinct colors are used to represent different lines. The T_3_ transgenic lines were analyzed using three biological replicates per line.

The VIP analysis revealed that the top 12 metabolites (out of 30) were identified as discriminating metabolites (VIP ≥ 1) between transgenic and non-transgenic tissues (P< 0.05). Significant higher contents of glucose-6-phosphate, glucose, betaine, mannose, citrate, N-carbomylaspartate, galactose, trans-4-hydroxy proline, glutamate, valine, methionine, and proline were upregulated by transgenic cowpea pods (SI Fig. 4).

To effectively compare the differences caused by cry1Ab expression, pairwise comparisons were performed using the supervised OPLS-DA approach. The random permutation test (100 times) on the OPLS-DA model confirmed the difference between control and transgenic cowpea (Fig. 5 A-E). The S-plot also confirmed the higher relative content of methionine, proline, galactose, threonine, fructose, valine, and glucose-6-phosphate in transgenic pods compared to control (Fig. 5 F-J).

**Figure 5:**
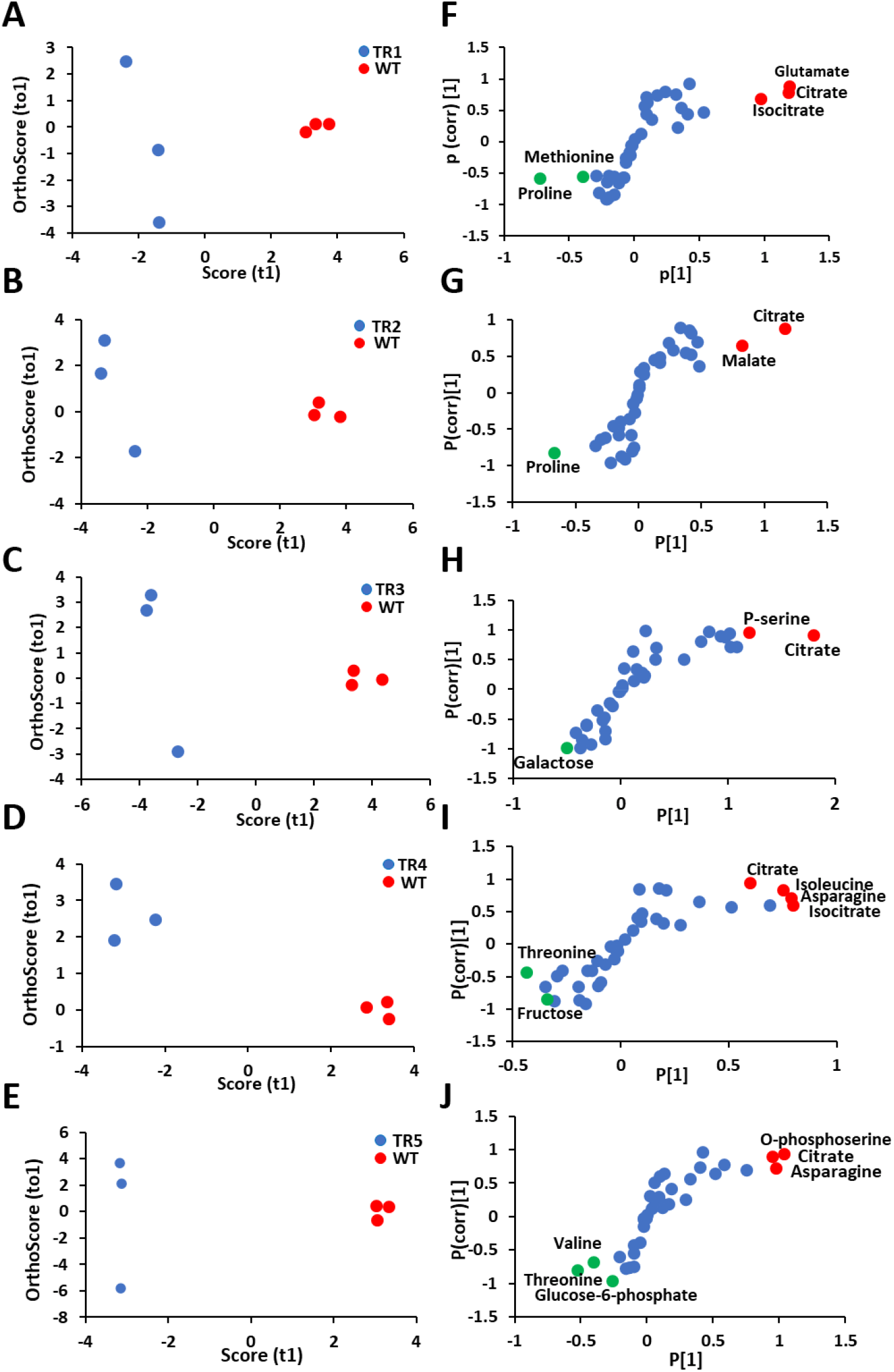
Statistical analysis of transgenic (line #5:TR1, line #12:TR2, line #17:TR3, line #30:TR4, and line #38:TR5) and non-transgenic (WT) samples. OPLS-DA score plot **(A-E)** and corresponding loadings S-plot **(F-J)** for discrimination of transgenic and non-transgenic cowpea pods. The T_3_ transgenic lines were analyzed using three biological replicates per line (n=3).

### Metabolic Pathway Analysis

The Web tool, MetaboAnalyst 5.0, was used to perform the biological pathway analysis. Metabolic pathway analysis (MetPA) suggested that thirteen pathways might be altered by the *cry1Ab* expression in cowpea based on the statistically significant value (p < 0.05) and impact factor > 0.1. This includes, (1) starch and sucrose metabolism, (2) alanine, aspartate and glutamate metabolism, (3) galactose metabolism, (4) valine, leucine and isoleucine biosynthesis, (5) cysteine and methionine metabolism, (6) glyoxylate and dicarboxylate metabolism, (7) arginine and proline metabolism, (8) amino sugar and nucleotide sugar metabolism, (9) glycine, serine and threonine metabolism, (10) arginine biosynthesis, and (11) citrate cycle (TCA cycle), (12) carbon fixation by Calvin cycle, (13) glutathione metabolism. The relationship between several metabolites and relevant metabolic pathways and the primary metabolites process is depicted in (Fig. 6 and Table 1).

**Figure 6.**
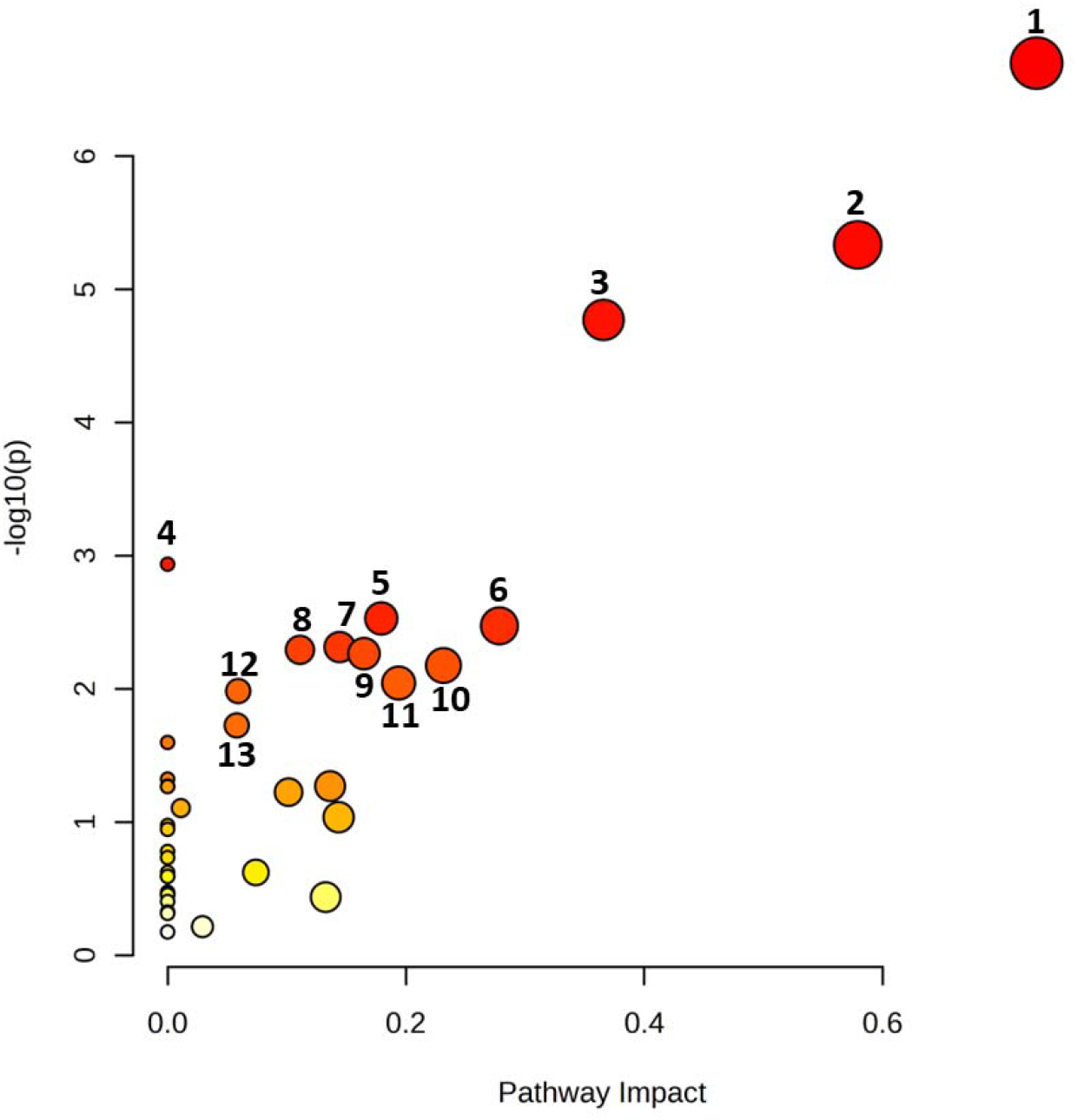
Metabolic pathways analysis from pods of transgenic cowpea lines expressing cry1Ab. Analysis carried out using MetaboAnalyst 5.0. Differences were considered statistically significant with P values⍰<⍰0.05 and an impact factor threshold >⍰0. Altered metabolic pathways of cry1Ab cowpea samples was observed. (1) Starch and sucrose metabolism, (2) Alanine, aspartate and glutamate metabolism, (3) Galactose metabolism, (4) Valine, leucine and isoleucine biosynthesis, (5) Cysteine and methionine metabolism, (6) Glyoxylate and dicarboxylate metabolism, (7) Arginine and proline metabolism, (8) Amino sugar and nucleotide sugar metabolism, (9) Glycine, serine and threonine metabolism, (10) Arginine biosynthesis, and (11) Citrate cycle (TCA cycle), (12) Carbon fixation by Calvin cycle, (13) Glutathione metabolism.

### Agronomic performance and segregational analysis

The agronomic traits of T_3_ transgenic plants expressing cry1Ab and their non-transgenic counterparts were assessed to determine if the incorporation of the Bt gene resulted in any phenotypic changes (Fig. 7). Morphologically, there were no observable differences between the transgenic and non-transgenic plants. Statistical analysis revealed no significant differences (P < 0.05) in key agronomic traits, including plant height, seed number per plant, seed length, and ten seed mass of transgenics were not different from non-transgenic plants (Fig. 7). Line #30 showed lower number of branches compared to other transgenics and non-transgenic plants. However, the pod number per plant was observed to be higher in line #30 and #38 compared to other plants (Fig. 7). Furthermore, transgenic plants exhibited no differences in growth or fertility compared to non-transgenic plants.

**Figure 7.**
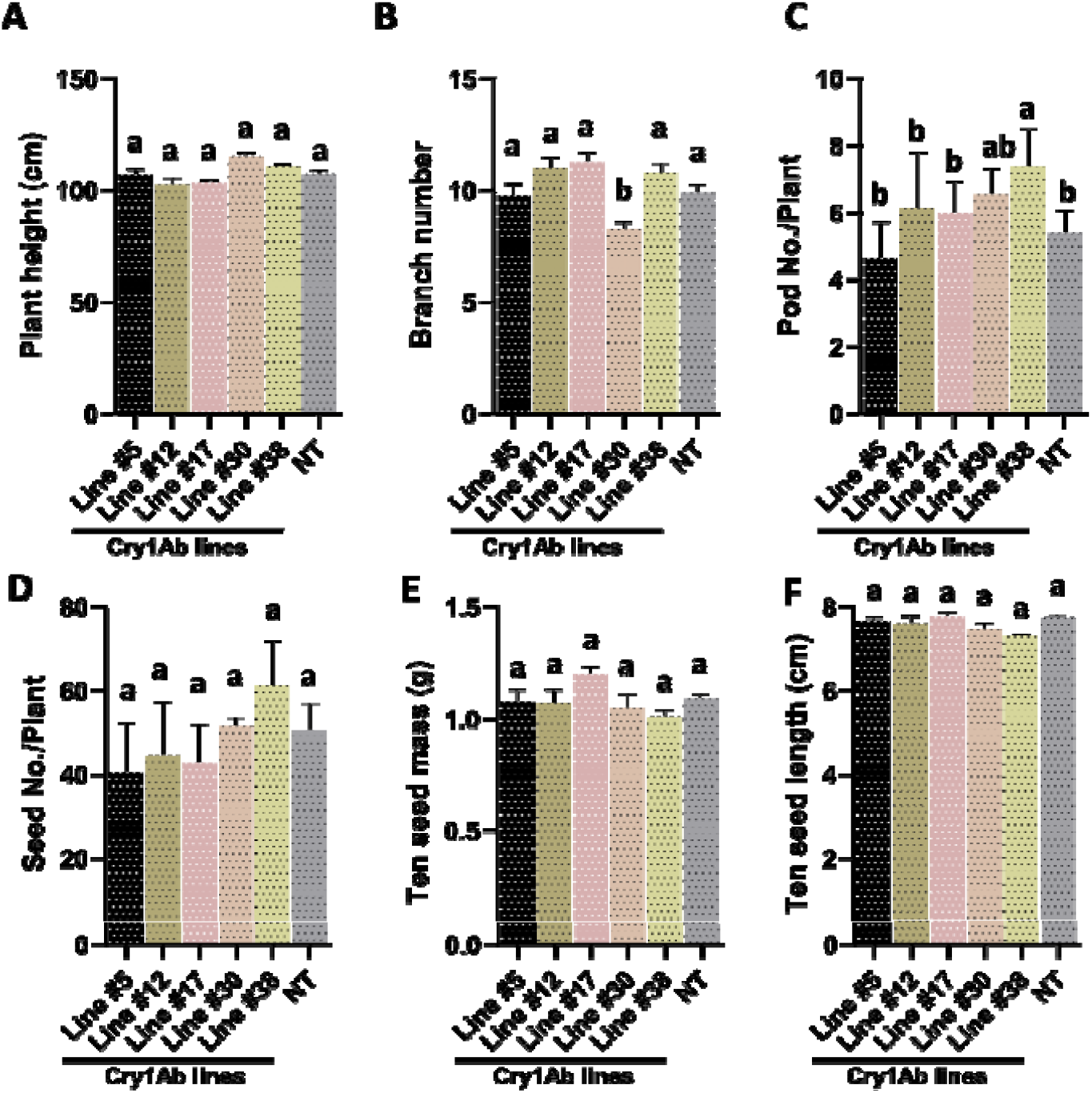
Evaluation of agronomical performance. Transgenic and non-transgenic cowpea lines were evaluated under field condition (poly-house). The physiological parameters including plant height **(A)**, branch number **(B)**, pod number/plant **(C)**, seed number/plant **(D)**, ten seed mass **(E)**, and ten seed length **(F)** were shown. Bars represent mean values ± standard deviation (SD) from n=3 biological replicates. Statistical significance was determined using one-way ANOVA followed by Tukey’s multiple comparison test in GraphPad Prism 8.0. Bars shares at least one common letter (a, b) are not significantly different from each other (P> 0.05), while bars labelled with different letters indicate statistically significant differences (P< 0.05).

## Discussion

Cowpeas are cost-effective protein-rich grain legumes critical for nutritional security and sustainable agriculture in developing countries. The legume pod borer (*M. vitrata*) is the most devastating insect pest of cowpea. Till date, no source of complete-resistance has been identified in the gene pool of cultivated cowpea. Amelioration of insect damage can add to cowpea production, reduce insecticide use and help in sustainable production (Muthuvel et al. 2021). Sources of resistance to the insect pest have not been identified in cowpea germplasm, which limited the approach of conventional breeding to develop resistant varieties(Horn and Shimelis 2020; Smith 2021). Genetic engineering of crops using genes encoding insecticidal crystal proteins or δ-endotoxins from B thuringiensis (Bt) has been shown to confer resistance to a variety of insects and pests in various crops (Huang 2021; Tilgam et al. 2021). Among others, Cry1Ab has been demonstrated to be highly toxic against early instars of *M. vitrata* (Srinivasan et al. 2021). Previous field studies have evidenced that cry1Ab provides effective protection for cowpea against *M. vitrata*. Artificial infestation of *M. vitrata* larvae, along with an integrated pest management strategy, resulted in nearly complete protection in transgenic cowpea lines expressing *cry1Ab* protein compared to non-transgenic cowpea lines (Nboyine et al. 2024). These findings suggest that *cry1Ab* could be a promising candidate for integrating insect resistance into cowpea.

We have developed transgenic cowpea lines expressing a codon-optimized cry1Ab gene from B. thuringiensis and verified their efficacy against the legume pod borer *M. vitrata* as well as *H. armigera*. Cowpea transformation protocol has been standardized and used to generate many transgenic lines to confer various biotic and abiotic stress tolerance in our lab (Kumar et al. 2017, Kumar et al. 2022). Following the same protocol, we have generated 65 putative transgenic cowpea lines. The transmission of the transgene/inheritance was confirmed by PCR analysis at each generation and molecular analyses was calculated to verify Mendelian segregation. Interestingly, a few transgenic events were found to be positive for the selectable marker *nptII* but negative for both *cry1Ab* and gus gene. This may be due to partial T-DNA integration or genomic rearrangements during T-DNA integration or chimerism in regenerated plants could result in such incomplete or tissue-specific transgene presence. Another possible explanation includes false positives during PCR screening, particularly if low-copy or truncated insertions occurred. These factors highlight the complexity of T-DNA integration and underscore the need for thorough molecular validation of transgenic events. We have observed few homozygous lines in T_2_ generation, based on PCR analysis. However, in most of the cases we have obtained mixed populations of homo and hemizygous lines T_2_ generation. The five homozygous PCR confirmed progenies with Mendelian segregation were used for characterization of the transgenic lines.

Southern blotting revealed single copy of the *cry1Ab* gene in the HindIII-digested genomic DNA of the selected transgenic cowpea lines when probed with a sequence specific to *cry1Ab*, indicating its stable integration in the cowpea genome. The presence of the expected hybridization signals with genomic DNA fragments (>4.2 kb) in the majority of the transformed plants showed that the probed genes *cry1Ab* remained intact upon integration into the cowpea genome (Figure 1 G). It is preferable to have low-copy (one to three) insertions of the transgene into the plant genome using A. tumefaciens, as they tend to remain stable over several generations. The transgenic cowpea plants have shown an independent pattern of transgene expression due to the complex and random integration of foreign genes in the host genome following Agrobacterium-mediated transformation. Therefore, the inheritance of foreign genes in transgenic plants may exhibit complex patterns for both single and multiple genes.

Given the poor expression levels of native Bt insecticidal proteins in higher eukaryotes (Jadhav et al. 2020; Li et al. 2022), it’s critical to ensure optimal expression of insecticidal proteins for effective control of targeted insects (Liu et al. 2020). The optimal expression of Bt genes in plants is controlled by various factors including codon preference, AT content, mRNA destabilization sequences and putative polyadenylation signals (Watts et al. 2021) (Perlak et al. 2001; Gatehouse 2008). Codon optimization has been previously employed in various crops, such as cotton (Zafar et al. 2022; Siddiqui et al. 2023) (Perlak et al. 1990), soybean (Fang et al. 2024) (Stewart et al. 1996), chickpea (Singh et al. 2022) (Das et al. 2017), and cowpea (Kumar et al. 2021) (Bett et al. 2017), to enhance expression levels. In our current study, we adopted a similar approach to maximize cry1Ab expression in transgenic cowpea. This involved increasing the GC content of the coding sequence and removing polyadenylation and mRNA destabilization sequences without altering the amino acid sequence. Additionally, codon usage was optimized to enhance translation in plants by incorporating plant-preferred codons. Relative fold expression by real-time PCR analysis of *cry1Ab* transcripts in transgenic cowpea plants, driven by the constitutive CaMV35S promoter, revealed higher expression levels of *cry1Ab* transcripts in both leaves and immature pods (Figure 1 E and F). In our transgenic lines, *cry1Ab* expression in leaves and immature pods reached 3 to 10-fold and 6 to 13-fold expression relative to ubiquitin, respectively. Notably, the expression of *cry1Ab* in these tissues surpasses that reported in previous studies on cowpea (Addae et al. 2020; Majumder et al. 2020; Eckerstorfer et al. 2022). Given the critical importance of achieving optimal expression of cry genes in field crops for effective pest control strategies, our transgenic lines emerge as potential candidates for integrating existing Cry lines into comprehensive field studies. Furthermore, our investigation revealed variations in cry1Ab expression across independent transgenic cowpea lines (Figure 1 E and F). These variations likely arise from factors such as positional effects, transgene copy numbers, and the choice of promoter for transgene expression. Notably, similar fluctuations in cry gene expression have been documented in numerous other crops engineered to confer resistance against lepidopteran insects (Siddiqui et al. 2023; Fang et al. 2024; Singh et al. 2022; Kumar et al. 2021). This underscores the importance of meticulous optimization and rigorous evaluation protocols in transgenic crop development. The results from PCR, Southern blotting, and RT-PCR analyses of transgenic cowpea plants decisively confirmed the stable integration without any rearrangements of the cry1Ab gene in transgenic plants and also in subsequent generations.

The effectiveness of selected cowpea lines expressing the Cry1Ab protein was assessed against MPB larvae by feeding them leaves and pods. Western blot analysis, followed by ELISA, confirmed stable cry gene expression. Among the T_3_ progenies, line #30 exhibited the highest Cry1Ab expression (7.6 µg/g in leaves and 8 µg/g in pods), while line #5 showed the lowest expression (7.6 µg/g in leaves and 8 µg/g in pods). The expression-based selection process successfully identified the high-expressing events in this research. Notably, the pods accumulated 6.6% more toxin on average than the leaves in the corresponding transgenic lines. Previous studies have documented the variability in Bt-endotoxin expression among transgenic plants driven by CaMV35S promoters. This variability has been attributed to factors such as the position effect of gene integration, surrounding flanking sequences, chromatin context, increased DNA methylation with plant age, and physiological changes affecting the stability of foreign proteins within plant tissues (To et al. 2021; Rurek and Smolibowski 2024). In contrast, our analysis of transgenic cowpea lines demonstrated that the expression of the Cry1Ab protein, as verified through Western blot and ELISA assays, remained robust and consistent in both leaves and immature pods across three successive generations. This stability highlights the potential for reliable expression of Bt-endotoxins in cowpea, minimizing the concerns raised in earlier observations.

The Bt-toxin expression levels observed in this study surpass those reported for other grain legumes (Acharjee and Higgins 2021; Singh et al. 2023) (Sanyal et al. 2005; Ramu et al. 2012; Singh et al. 2023) and represent the highest recorded Cry protein levels in cowpea (Addae et al. 2020; Kumar et al. 2021). This higher expression may be attributed to the use of a codon-optimized cry1Ab gene, specifically engineered to enhance mRNA stability, eliminate polyadenylation sites and splicing sequences, and optimize ATG consensus flanking nucleotides for efficient translation initiation. Additionally, the transgene’s integration into a transcriptionally active region of the host genome likely contributed to this robust expression (Watts et al. 2021) (Perlak et al. 1990; Sardana et al. 1996). Codon optimization has been extensively employed in Bt-cry genes to enhance insect resistance across various plant species (Ferry and Gatehouse 2010). High-level expression of codon-optimized *cry1Ac, cry1Ab*, and *cry1C* genes has been previously demonstrated in transgenic cotton, tomato, and tobacco, respectively (Zafar et al. 2022; Siddiqui et al. 2023; Fernandes et al. 2023; Wang et al. 2024) (Perlak et al. 1990).

Furthermore, increased expression of Cry2Aa in cowpea has been shown to improve resistance against lepidopteran pests (Singh et al. 2018; Kumar et al. 2021).

The expression levels of a transgene are significantly influenced by the choice of promoter driving its transcription and translation into functional protein. Historically, the CaMV35S promoter has been widely used to achieve constitutive expression of cry1Ab in transgenic cowpea, and the first pod-borer-resistant cowpea released for commercial use in Nigeria utilized the CaMV35S promoter for gene regulation (https://aaccnet.confex.com/aaccnet/2020/meetingapp.cgi/Paper/5721). However, recent studies have shown that the green tissue-specific RuBisCO small subunit (rbcS) promoter can drive Cry1Ac expression to levels 1.5 times higher than those achieved with CaMV35S and Ubi promoters in chickpea (Boruah et al. 2023; Hazarika et al. 2021). These findings suggest that future research should explore the use of the rbcS promoter to achieve enhanced endotoxin protection through higher transgene expression.

Our study adds to the expanding body of research focused on the effectiveness of transgenic plants expressing insecticidal proteins in controlling major lepidopteran pests. Through in vitro insect feeding bioassays using larvae of *M. vitrata* and H. armigera, we observed a marked increase in insect-induced damage in non-transgenic control compared to transgenics. This was directly linked to the presence of *Cry1Ab* toxins in the leaves and immature pods of the transgenic plants. Notably, all five transgenic lines exhibited resistance to insect damage, with some variation in effectiveness, likely due to differences in *cry1Ab* gene expression levels. Our results are consistent with previous studies on the role of transgenic plants in pest management. For instance, cowpea lines expressing the Cry2Aa protein showed substantial protection against Maruca pod borer larvae, with mortality rates exceeding 90% (Kumar et al. 2021). Similarly, chickpea plants co-expressing cry1Ab and cry1Ac genes exhibited enhanced toxicity, providing broader protection against pests such as H. armigera (Koul et al. 2022). Some Bt-transgenic chickpea lines even achieved 100% mortality, underscoring the potential of Bt-transgenics as an effective pest management tool (Hazarika et al. 2021).

Moreover, our *cry1Ab* cowpea transgenics demonstrated enhanced insecticidal efficacy not only against *M. vitrata* but also against H. armigera. Interestingly, although Cry1Ab protein levels were lower in the leaves compared to pods of transgenic cowpea lines, higher H. armigera mortality was observed in leaf-feeding assays. This apparent contradiction can be attributed to several factors. Firstly, H. armigera larvae show a feeding preference for softer leaf tissues during early instars, leading to greater ingestion of Cry1Ab despite lower expression levels in leaves (Zalucki et al. 1994; Sharma et al., 2005). Secondly, Cry1Ab may be more evenly distributed or bioavailable in leaf tissues, whereas physical structures in pods may hinder effective ingestion. Additionally, plant-derived secondary metabolites such as flavones, commonly found in leaves, may interact synergistically with Cry toxins to enhance their toxicity (Wang et al., 2021). Furthermore, recent studies suggest that Cry1Ab’s toxicity depends on binding to specific receptors in the insect midgut, including prohibitin and cadherin-like proteins, whose expression or accessibility could vary depending on the tissue consumed (Sena da Silva et al., 2021). Therefore, the observed mortality is likely influenced not just by absolute Cry1Ab levels, but also by tissue-specific feeding dynamics, toxin accessibility, plant metabolite interactions, and receptor-mediated mechanisms. The observed reduction in larval mortality in transgenic cowpea compared to larvae fed on non-transgenic control plants confirms the effective expression of the toxin protein in these transgenic lines. These findings provide a strong foundation for future crop protection strategies, suggesting that integrating Cry1Ab proteins into cowpea could be a promising approach to controlling *M. vitrata* and H. armigera infestations, thereby minimizing crop damage and enhancing agricultural sustainability.

Metabolomics has emerged as a powerful tool for providing deep insights into crop biology. The information obtained from metabolomic analyses can be effectively utilized in assessing phenotypic changes, identifying biomarkers, and tracking gene expression, while also enhancing the interpretation of other genomic data (Makhumbila et al. 2022). NMR metabolomics, in particular, is highly effective in studying plant defenses, both constitutive and induced, against biotic stressors(Mascellani Bergo et al. 2024). Our study revealed that the majority of metabolites in transgenic cowpea were upregulated, including amino acids, sugars, and other key metabolites (Fig. 6). Notably, carbohydrates such as sucrose and fructose were significantly more abundant in transgenics compared to non-transgenic cowpea. These results align with those of (Chang et al. 2012), who reported elevated levels of sucrose, mannitol, and glutamic acid in transgenic rice expressing *cry1Ac* and sck genes. Metabolic pathway analysis further highlighted the upregulation of the amino acid metabolism pathway, TCA cycle, and carbohydrate metabolism pathway (Fig. 6; Table 1). Our findings indicate that while no new metabolites were detected, there was a significant alteration in the abundance of existing ones. Similar outcomes were observed in maize transgenics overexpressing cry proteins (Liu et al. 2021; Liu et al. 2023). The use of both targeted and untargeted NMR metabolomics was crucial in evaluating the metabolite profile variations in cowpea leaves and pods resulting from transgene expression.

## Conclusion

In conclusion, our study demonstrated that the synthetic *cry1Ab* gene is a valuable strategy for enhancing MPB resistance. This is the first report of a transgenic cowpea variety using a synthetic *cry1Ab* gene, which also conferred resistance to H. armigera. The *cry1Ab* gene is a strong candidate for co-expression with other cry genes in developing Bt cowpea resistant to multiple insects, including MPB and H. armigera. Such combinations could delay resistance evolution, provide broad-spectrum protection, increase yields, boost the income of farmers, and reduce pesticide use. Our transgenic cowpea showed high insect mortality and holds promise for future genetic improvement programs, advancing durable insect resistance and sustainable agriculture.

## Author Contribution Statement

LS conceived and designed research, contributed new reagents/analytical tools and corrected the manuscript. MJ designed, conducted experiments and analyzed data and wrote the manuscript. DKM and SK assisted MJ in some experiments. IA provided the synthetic *cry1Ab* construct. VK conducted insect bioassay. All authors read and approved the manuscript.

---

**SI Figure 1:**
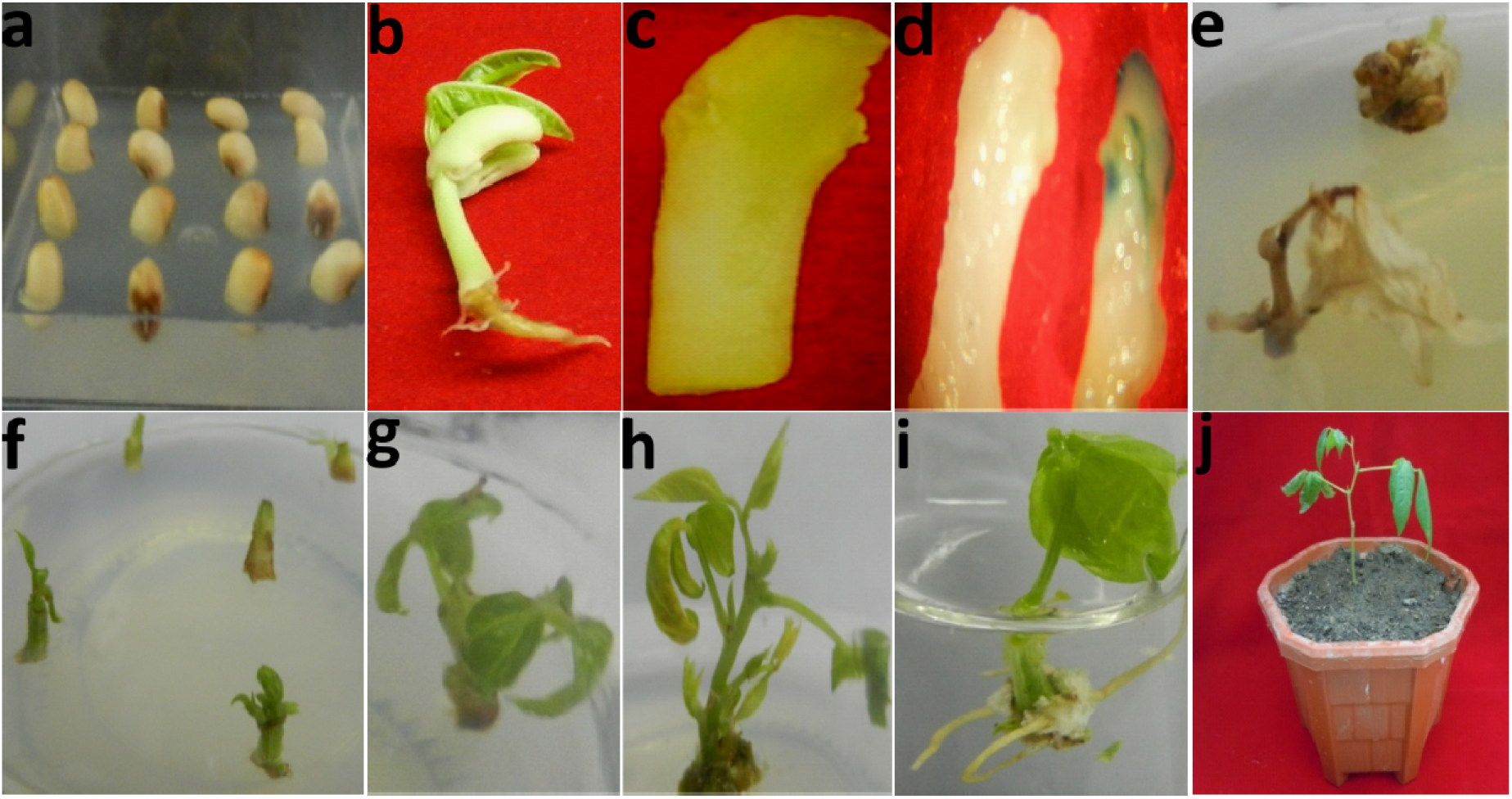
Stages of cowpea (*Vigna unguiculata*) cv. PUSA KOMAL through *Agrobacterium*-mediated transformation. a. Cowpea seeds in germination media b. Four days old cowpea seedling c. Cotyledonary node explants at the time of culture d. Transient gus expression in control (left) and transformed cotyledonary node (right) explants after 3 days of co-cultivation e. Control explants cultured on selection media f. Selection of transformed explants on selection media g. Shoot proliferation from explant in subsequent subculture h. Elongation of shoot within 4 week of culture i. In vitro rooting of selected shoot j. Plant hardened and maintained in transgenic green house.

**SI Figure 2.**
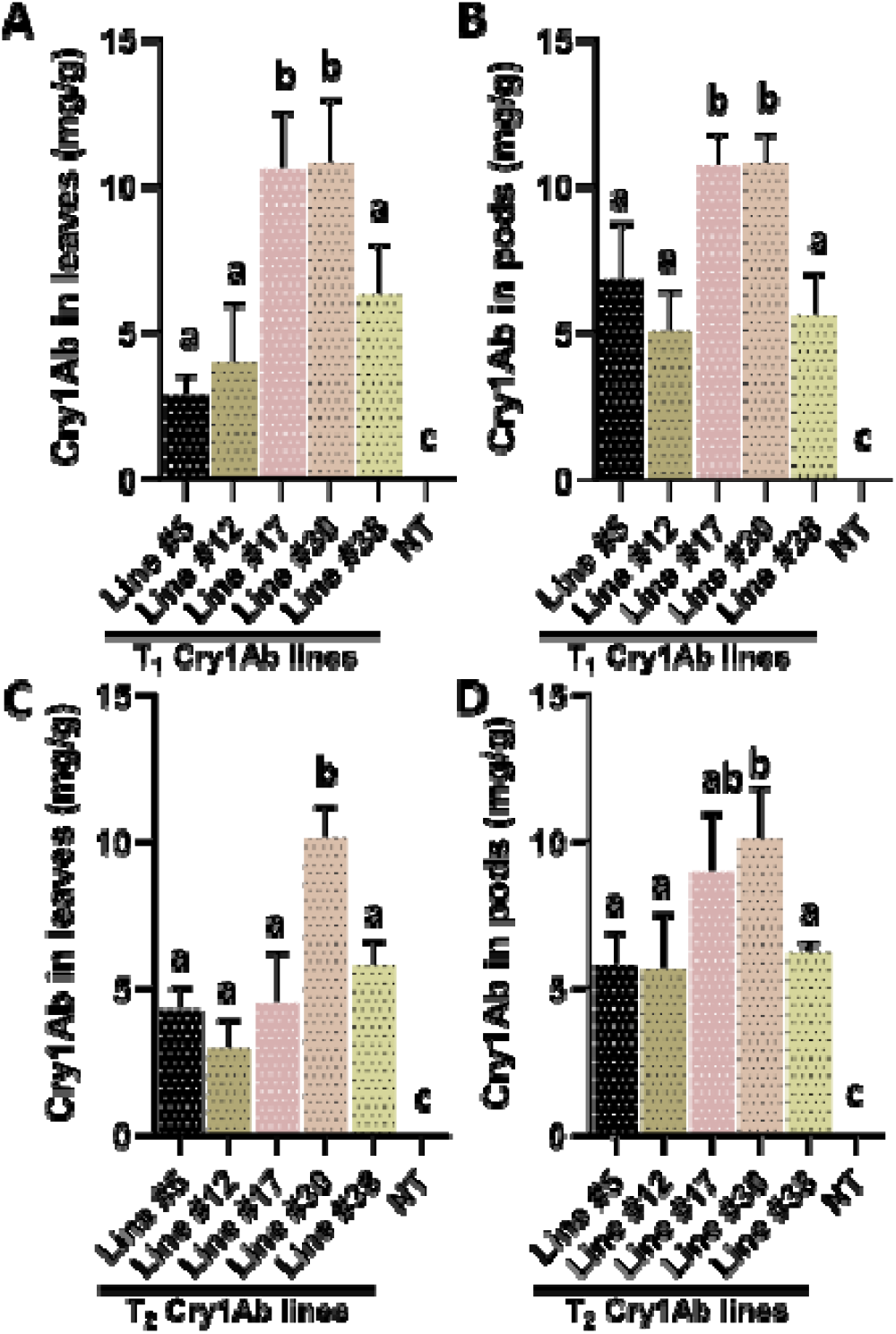
Quantitative estimation of Cry1Ab protein in T_1_ and T_2_ transgenic leaves (Line #5-38) **(A, C)**, pods (Line #5-38) and non-transgenics (NT) **(B, D)** using Cry1Ab-enzyme conjugate. Average quantity of total Bt-Cry protein of the transgenics is represented as mg g^−1^ TSP. Bars represent mean values ± standard deviation (SD) from n=3 biological replicates. Statistical significance was determined using one-way ANOVA followed by Tukey’s multiple comparison test in GraphPad Prism 8.0. Bars shares at least one common letter (a, b and c) are not significantly different from each other (P> 0.05), while bars labelled with different letters indicate statistically significant differences (P< 0.05).

**SI Figure 3.**
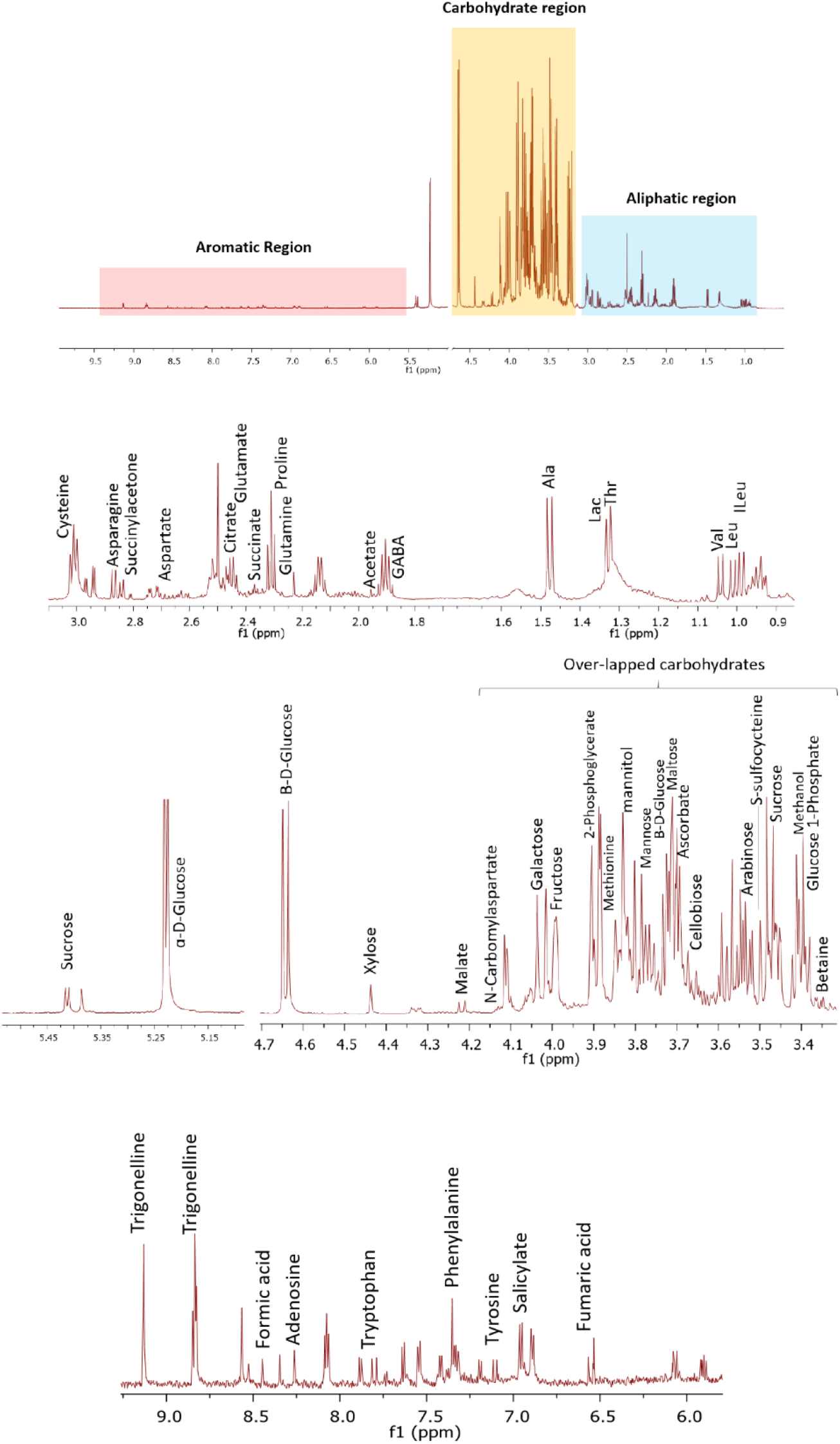
Representative ^1^H-NMR spectra obtained at 600 MHz from cowpea pods expressing *cry1Ab* are provided. The Aliphatic, carbohydrate, and aromatic regions within the spectra are highlighted. The individual compounds present in these regions are marked and shown.

**SI Figure 4:**
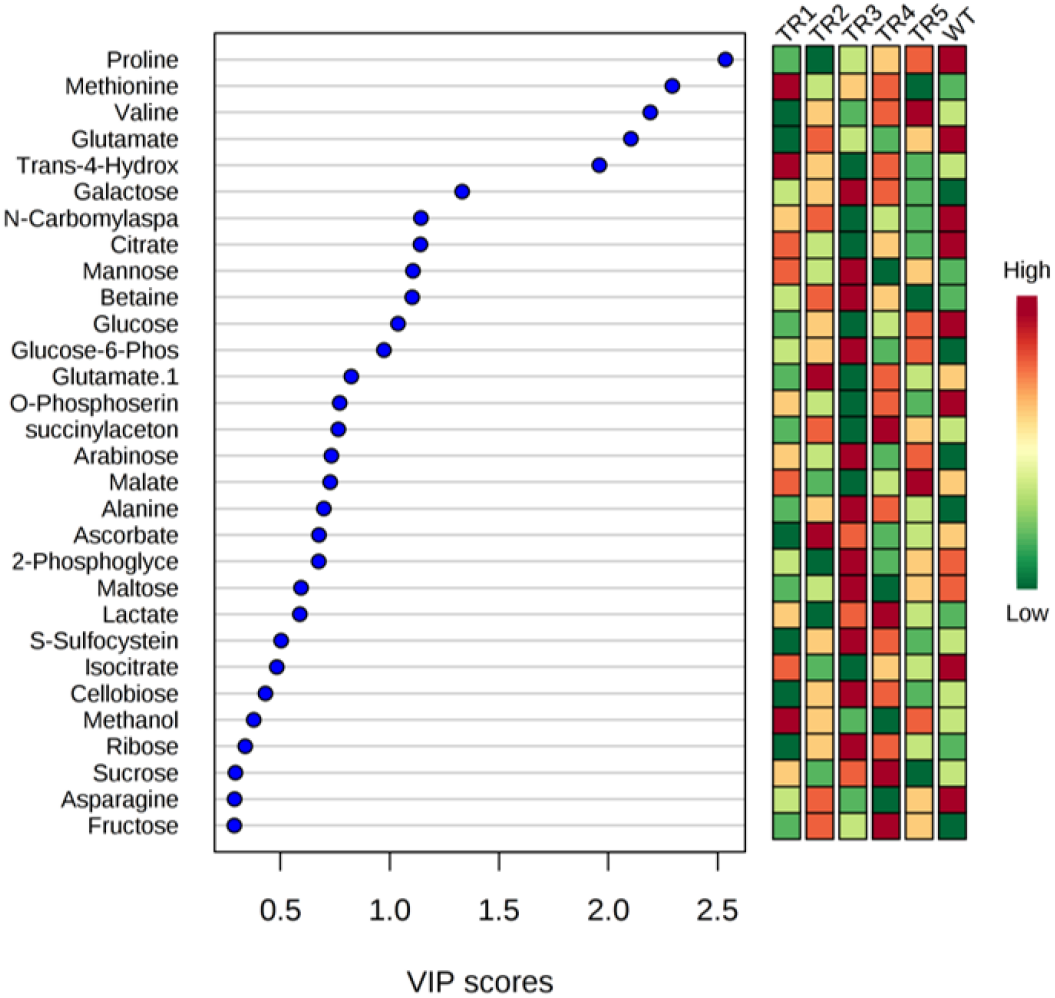
Top 30 metabolites (variables) present in transgenics (line #5:TR1, line #12:TR2, line #17:TR3, line #30:TR4, and line #38:TR5) and non-transgenic (WT) based on VIP score.

**SI Table 1.**
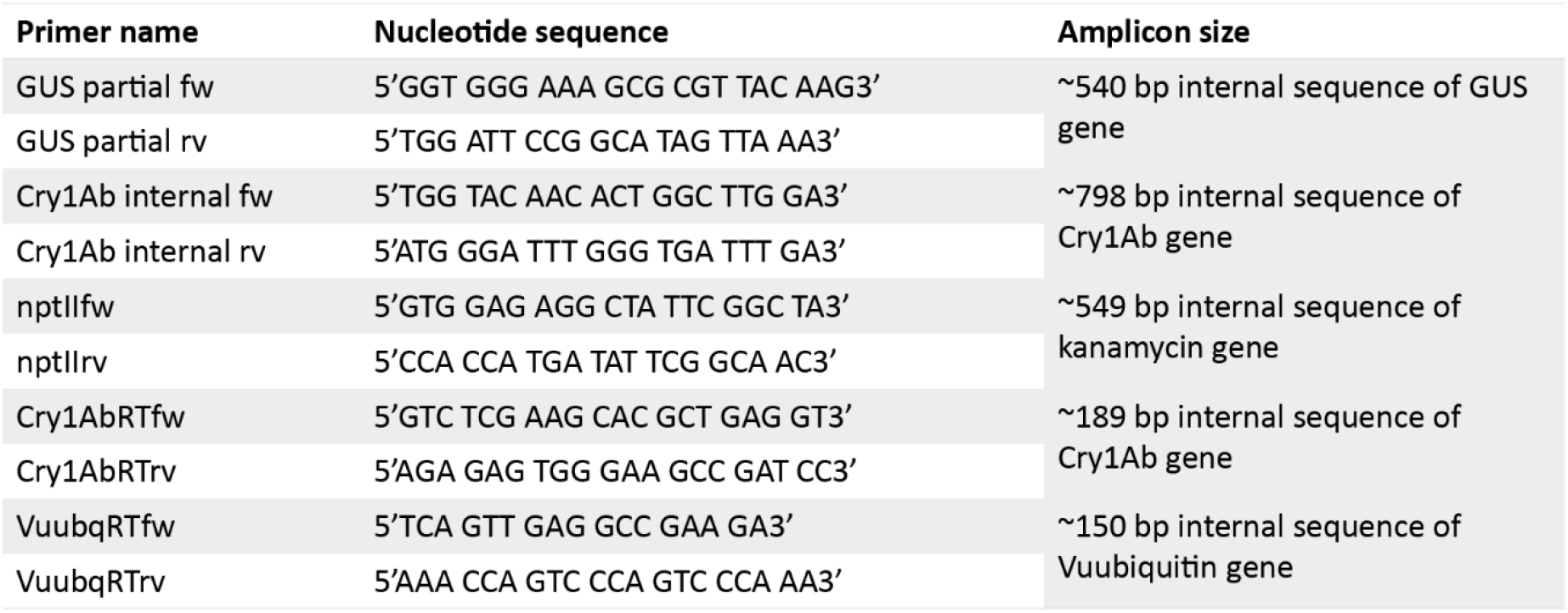
Primers used in our study.

**SI Table 2:** Identification of metabolites in the pods of both transgenic and non-transgenic cowpea expressing *cry1Ab* with their chemical shift and multiplicity.

| SN | Metabolites | Chemical shifts (ppm) | Multiplicity |
| --- | --- | --- | --- |
| 1 | Leucine | 0.96 | d |
| 2 | Valine | 0.99 | d |
| 3 | Isoleucine | 1.01 | m |
| 4 | Lactate | 1.33 | d |
| 5 | Threonine | 1.42 | dd |
| 6 | Alanine | 1.47 | d |
| 7 | Arginine | 1.91 | dd |
| 8 | Glutamate | 2.08 | m |
| 9 | Glutamine | 2.12 | m |
| 10 | Proline | 3.33 | m |
| 11 | Methionine | 3.99 | s |
| 12 | 4-Aminobutyrate | 3.01 | d |
| 13 | succinylacetone | 3.85 | s |
| 14 | Isocitrate | 4.05 | s |
| 15 | N-Carbomylaspartate | 4.22 | d |
| 16 | Trans-4-Hydroxyl-L-Proline | 3.47 | dd |
| 17 | Citrate | 2.49 | d |
| 18 | Asparagine | 2.94 | dd |
| 19 | Aspartate | 3.89 | dd |
| 20 | Malate | 4.34 | d |
| 21 | Cysteine | 3.98 | m |
| 22 | Xylose | 3.52 | m |
| 23 | Glucose | 4.64 | m |
| 24 | Maltose | 4.65 | m |
| 25 | Methanol | 3.35 | s |
| 26 | Mannose | 3.57 | m |
| 27 | Glucose-1-Phosphate | 3.36 | m |
| 28 | Cellobiose | 3.96 | m |
| 29 | S-Sulfocysteine | 4.17 | dd |
| 30 | Galactose | 3.98 | m |
| 31 | Glucose-6-Phosphate | 3.6 | m |
| 32 | Arabinose | 3.78 | m |
| 33 | Fructose | 4.05 | m |
| 34 | Ribose | 4.21 | m |
| 35 | Mannitol | 3.86 | dd |
| 36 | Sucrose | 5.41 | d |
| 37 | Ascorbate | 4.01 | m |
| 38 | 2-Phosphoglycerate | 4.47 | m |
| 39 | Betaine | 3.27 | s |
| 40 | O-Phosphoserine | 4.18 | m |

